# Proteomic and *N*-glycomic comparison of synthetic and bovine whey proteins and their effect on human gut microbiomes

**DOI:** 10.1101/2024.09.18.613515

**Authors:** Matthew Bolino, Hatice Duman, İzzet Avcı, Hacı Mehmet Kayili, Juli Petereit, Chandler Zundel, Bekir Salih, Sercan Karav, Steven A. Frese

## Abstract

Advances in food production systems and customer acceptance have led to the commercial launch of dietary proteins produced via modern biotechnological approaches as alternatives to traditional agricultural sources. At the same time, a deeper understanding of how dietary components interact with the gut microbiome has highlighted the importance of understanding the nuances underpinning diet-microbiome interactions. Novel food proteins with distinct post-translational modifications resulting from their respective production systems have not been characterized, nor how they may differ from their traditionally produced counterparts. To address this, we have characterized the protein composition and *N*-glycome of a yeast-synthesized whey protein ingredient isolated from commercially available ice cream and compared this novel ingredient to whey protein powder isolate derived from bovine milk. We found that despite strong similarities in protein composition, the *N*-glycome significantly differs between these protein sources, reflecting the biosynthetic machinery of the production systems. Further, the composition profile and diversity of proteins found in the synthetic whey protein were lower relative to bovine whey protein, despite both being predominantly composed of β-lactoglobulin. Finally, to understand whether these differences in *N-*glycome profiles affected the human gut microbiome, we tested these proteins in an *in vitro* fecal fermentation model. We found that the two whey protein sources generated significant differences among three distinct microbial compositions, which we hypothesize is a product of differences in *N*-glycan composition and degradation by these representative microbial communities. This work highlights the need to understand how differences in novel biotechnological systems affect the bioactivity of these proteins, and how these differences impact the human gut microbiome.

## 1. Introduction

There has been considerable interest in the development of novel protein sources to meet increasing global demands for dietary protein, especially those that meet consumer interests related to animal welfare, CO_2_ emissions, and land use practices (1). The interest in novel protein sources has resulted in significant commercial investment in developing viable protein sources that mimic or replace animal derived food products. For example, popular plant-based milk alternatives lack many of the sensory and nutritional qualities of bovine milk (2,3) and clinical recommendations caution against the substitution of plant-based milk alternatives for infants and children (4). Still, there has been a growing interest in developing a more robust alternative for bovine milk that is both functional and nutritionally more complete than currently available milk alternatives (5).

Recent developments in biosynthesis, fermentation, and heterologous gene expression technologies have resulted in the potential to produce a wide variety of dietary ingredients including milk oligosaccharides (6), lipids (7), and protein (8). Biotechnological advances have also enabled the complex synthesis of large organic molecules and feedstocks for further organic molecule production (9,10). More recently, this technology has been applied to the development of dietary ingredients derived from genes originating from other organisms to produce food proteins *in vitro*, the result of which is now commercially available in the United States (11).

While these novel technologies and ingredients may be conceptually attractive alternatives for consumers, there is little research characterizing these replacements for traditional food ingredients. For some novel food ingredients, such as complex human milk oligosaccharides or other synthetic organic chemical feedstocks, the chemical identity can be compared directly. However, complex biomolecules such as proteins are subject to post-translational modifications including *O-* and *N*-linked glycosylation, which requires additional characterization to describe the structure of dietary proteins. These post-translational modifications have been the subject of intense research in recent years, which has shown that bioactive function, protein functionality, and interactions with the microbiome are affected by protein glycosylation or glycation (12–15).

Importantly, milk is a complex biological fluid that provides a sole source of nutrition for developing mammals and contains a wide variety of proteins, lipids, carbohydrates, and many bioactive compounds (16). The composition of milk has evolved to nourish, protect, and shape the development of the growing infant, and a full reconstruction of this nutritional and bioactive milieu is currently not feasible (11). However, where milk is a functional ingredient in other foods, it is possible that the primary organoleptic functions may be reconstructed without the associated complexity of the naturally occurring bioactive fluid. In the case of whey protein, whey is typically derived from cheese manufacturing which retains milk caseins as the coagulated protein that forms cheese while the whey proteins are removed during production and dried as a distinct ingredient. Compared to milk, whey is compositionally less complex and is predominantly made up of only a limited number of proteins, making it a more tractable system to replicate (11,17).

Here, we examine the proteome and *N*-glycome of a commercially available synthetic whey protein ingredient and compare it to bovine-derived whey protein isolate to determine how both protein identity and glycosylation patterns differ as a result of their respective biosynthetic origins. Using mass spectrometry to characterize these protein samples, we were able to determine the protein composition and how *N*-glycosylation motifs differ when produced in a yeast host, where native *N-*glycosylation synthesis is distinct from mammalian *N*-glycosylation systems, and how these differences affect the human gut microbiome using an *in vitro* model system.

## 2. Methods

### 2.1 Protein Purification

Synthetic whey proteins were purified from commercially available, milk protein-based foods advertised to contain these proteins, purchased at a Reno, NV market. Samples were centrifuged at 4000RPM for 20 mins to separate fat and centrifuge particulates. The de-fatted solution was then subjected to 4 rounds of ethanol precipitation by adding 4 volumes of ice-cold ethanol, incubation at -20°C overnight, then followed by centrifugation at 4C (4000RPM, 25 mins) to remove residual sugars and other remaining contaminants. Bovine whey protein samples were obtained from commercial, powdered whey protein isolate and purified in the same manner after suspension in water (20% w/v). The protein samples were subsequently aliquoted and dried at 30°C in a vacuum centrifuge. Purified protein was quantified using a Qubit BR Protein assay (ThermoFisher Scientific, Waltham, MA USA) and then evaluated via denaturing SDS-PAGE in a 4-15% acrylamide gel, stained with Coomassie (Bio-Safe^™^Coomassie; Bio-Rad Laboratories, Inc, Hercules, CA USA).

### 2.2 Proteomic analysis

Purified protein extracts (n = 5 per protein source) were reduced, alkylated and digested with a trypsin/Lys-C protease mixture using Thermo Scientific EasyPep Mini MS Sample prep kit (Cat #A40006). For LC-MS, peptides were trapped prior to separation on a 300 μm i.d. x 5 mm C18 PepMap 100 trap (Thermo Scientific, San Jose, CA) and separated on a 50 cm uPAC C18 nano-LC column (PharmaFluidics, Ghent, Belgium) with a 15 μm tip using an UltiMate 3000 RSLCnano system (Thermo Scientific, San Jose, CA). Mass spectral analysis was performed using an Orbitrap Eclipse mass spectrometer (Thermo Scientific, San Jose, CA) using data-independent acquisition (DIA). Six gas phase fractions (GPF) of the biological sample pool were used to generate a reference library. The GPF acquisition used 4 m/z precursor isolation windows in a staggered pattern (GPF1 398.4-502.5 m/z, GPF2 498.5-602.5 m/z, GPF3 598.5-702.6 m/z, GPF4 698.6-802.6 m/z, GPF5 798.6-902.7 m/z, GPF6 898.7-1002.7 m/z). Samples were analyzed on an identical gradient as the GPFs using a staggered 8 m/z window scheme over a mass range of 385-1015 m/z. Library generation and data analysis were performed using Spectronaut software (Biognosys, Schlieren, Switzerland) and peptide mapping was repeated against a protein database that included all known bovine milk proteins (https://www.dallaslab.org/resources). Peptides mapping to porcine trypsin and a serine protease from *Achromabacter* were omitted from analysis as they were added during sample preparation.

### 2.3 N-glycan profiling

Enzymatic deglycosylation of denatured protein samples was performed with PNGase F (1 Unit/μL) obtained from Promega (Madison, WI, USA). First, dried purified protein samples (1 mg) were dissolved in 50 μL of 2% SDS and denatured by incubations at 60°C. Denatured protein samples were then mixed with 2% NP-40 solution and 5X PBS and 1 U PNGase F was added and incubated at 37°C overnight. Finally, the samples were centrifuged, and supernatants were collected for further analysis. After enzymatic deglycosylation of protein samples, released *N*-glycans from each sample were labeled with a 2-AA tag. 50 μL of 2-AA tag (48 mg/mL^−1^ in DMSO/glacial acetic acid, 7/3, v/v) and 50 μL of 2-sodium cyanoborohydride (60 mg/mL in DMSO/glacial acetic acid, 7:3 w/v) were added to the released glycan samples (50 μL). Subsequently, the mixtures were incubated at 65°C for two hours. Purification of *N*-glycans was achieved by solid-phase extraction cartridges containing cellulose and porous graphitized carbon materials, as previously described (18). MALDI-TOF MS analysis of 2-AA labeled *N*-glycans from bovine and synthetic whey protein samples was carried out on a Bruker rapifleX™ MALDI Tissuetyper™ (Bruker Daltonik GmbH, Bremen, Germany) equipped with a SmartBeam 3D laser system. On the AnchorChip MALDI-target plate, the purified *N*-glycans (1 μL) were spotted and allowed to dry. Then, 1 μL of DHB matrix (5 mg/mL^−1^ in ACN/H_2_O, 1/1, v/v comprising 0.1% ortho-phosphoric acid) was added. The analysis included a 20 kV acceleration voltage, a 160 ns extraction delay, and the summation of 8000 shots at 2000 Hz for each spectrum. The mass range of 1000-5000 Da was used to produce all spectra using a random walk pattern in negative ion and reflectron mode. Data obtained by MALDI-TOF MS analysis were processed using Flex Analysis v.4.0 software (Bruker Daltonik Gmbh). Peaks of 2-AA labeled *N*-glycans were inserted into ProteinScape software including the GlycoQuest algorithm (Bruker Daltonik GmbH, Bremen, Germany) for glycan identification. Total area normalization was used to determine the relative abundance of individual *N*-glycans (mass-intensity based). All experiments were performed with three technical replicates.

### 2.4 Fecal sample collection and characterization

Fecal samples were collected from healthy individuals under supervision of the University of Nevada, Reno Institutional Review Board (Approval #1751022), from which multiple aliquots were collected and stored at -80°C. DNA was extracted from these samples as previously described (19) using a ZymoBiomics DNA Miniprep kit (Zymo Research, Irvine, CA USA) according to the manufacturer’s instructions, which included five rounds of bead beating for one minute, followed by incubation on ice for one minute. The resulting DNA was subjected to 16S rRNA sequencing of the V4 region using a previously described dual-indexed barcoding strategy (20) with recent modifications to the amplification sequences (21,22). Amplicons were generated in a HEPA-filtered laminar flow cabinet dedicated to PCR preparation and decontaminated before and after use. Kit and reaction controls were also included in downstream sequencing. Reactions were carried out using 200 nM of each primer, 0.5 mM added MgCl2, and GoTaq Master Mix (Promega; Madison, WI, USA) in 25 μL volumes with the following program in a MJ Research PTC-200 thermocycler: 94°C for 3 min, 25 cycles of 94°C for 45 s, 50°C for 60 s, and 72°C for 90 s, followed by a final extension at 72°C for 10 min. PCR reactions were pooled and purified with the High Pure PCR product purification kit (Roche Diagnostics; Mannheim, Germany) and 250bp paired-end sequencing was performed on an Illumina MiSeq at the Idaho State University Molecular Research Core Facility.

Demultiplexed sequencing data was analyzed with qiime2 (23). Reads were demultiplexed, trimmed to 200 bp, and joined, denoised and assigned to ASVs using DADA2 (24). Representative sequences were aligned with FastTree (25), and taxonomic assignments were assigned using the Silva database (v138, 99%; 24) with a feature classifier trained against the representative sequences. Distinct mircobial communities were determined using the genus-level classification method described by Arumugan et al (27) and available at https://enterotype.embl.de and in the supplementary data for this manuscript. Samples were rarefied to 2,000 reads per sample for diversity analyses, after determination by rarefaction analysis. ANCOM-BC (28) was used to identify differentially abundant taxa as noted.

### 2.5 In vitro batch fermentation and analysis

Six aliquots of fecal samples that could be assigned to one of three clusters, representing distinct human gut microbiota communities, identified among the total collection of fecal samples were diluted 1:10 (w/v) in cold sterile phosphate buffered saline containing 15% glycerol to maintain viability until *in vitro* fermentation. The six fecal samples within each of the three distinct clusters were then pooled in equal volumes to generate pooled and standardized inocula representative of each distinct microbial community. These three standardized inocula were stored at -80°C until use. Using an approach adapted from Aranda-Díaz et al., (29) and reported previously (30), each pooled community was inoculated (1% v/v) into 1 mL of a modified BHI medium in a deep-well 96-well plate as a control, or the same medium supplemented with bovine whey protein isolate (2% w/v) or fungal whey protein (2% w/v) with four independent replicates per treatment per plate. The anaerobic culture medium was composed of BHI supplemented with 3.5g/L of soluble starch, 0.3 g/L of L-cysteine HCl, 0.3 g/L of sodium thioglycolate, 1.5 mg/L of vitamin K1 and 0.3 mg/L of hemin and made anaerobic through mixed gas exchange in a Coy anaerobic chamber prior to fermentation. At inoculation, 100 μL of this culture was transferred to a 96-well plate and incubated at 37°C in a BioTek Epoch 2 96-well plate reader (Agilent Technologies; Santa Clara, CA) housed within an anaerobic chamber with one minute of shaking prior to an OD600nm measurement every 30 minutes for 24 hours. The deep-well plate was sealed with sterile film allowing for gas exchange and incubated at 37°C in the same anaerobic chamber. The anaerobic chamber is made anaerobic using three vacuum cycles of a mixed gas composed of 5% H_2_, 5% CO_2_, 90% N_2_. An oxygen indicator measuring oxygen in Parts Per Million (PPM) is housed within the anaerobic chamber to confirm oxygen-free conditions. After 12 and 24 hours, a replicate deep well plate was removed from the anaerobic chamber, centrifuged at 4°C for 30 mins (4,000 RPM), the supernatant was removed from the cell pellet into a new deep well plate and stored at -80°C. DNA was extracted from pelleted cells using a ZymoBiomics 96 DNA kit (Zymo Research, Irvine, CA USA) and subjected to 16S rRNA sequencing, as described above.

### 2.6 Statistical methods

Statistical tests were performed in R (v. 4.2.2) (31). Protein abundances were compared between groups using a nonparametric Kruskal Wallis test (32) using *ggpubr* (v. 0.4.0) and *rstatix* (v. 0.7.0) R packages (33,34). The protein and *N*-glycan abundances were used to calculate the Bray-Curtis distance (35) between respective sample types using adonis in the *vegan* R package (v. 2.6.4) (36). Fecal community composition and abundance was assessed using the weighted UniFrac distance and compared using adonis and PERMANOVA tests in qiime2 (v.2024.5). Figures were visualized using the *tidyverse, ggplot2* (v. 3.4.0), *ggpubr* (v. 0.4.0), and *viridis* (v. 0.6.2) R packages (33,37–39).

## 3. Results

### 3.1 Proteomic analysis and comparison to bovine whey

Mapping identified peptides to either an unrestricted protein database or a bovine-milk specific database produced a limited number of identified peptides in the case of both bovine whey and the synthetic whey product. Among the most abundant proteins identified through the unrestricted protein database (>1% relative abundance), β-lactoglobulin, α-lactalbumin, albumin, and casein S1 were the most common milk proteins to which peptides from bovine whey protein isolate could be mapped. Additional proteins identified included keratin proteins (KRT1, KRT2, KRT9, and KRT10), though at lower relative abundance (Table S1). When peptides from the bovine whey protein isolate were mapped to only bovine milk proteins, the most abundant proteins (>1% relative abundance) to which peptides were mapped were β-lactoglobulin, α-lactalbumin, albumin and casein S2. Additionally, we identified GLYCAM1 (glycosylation-dependent cell adhesion molecule 1) and lactadherin in this protein source (Figure 1, Table S1).

**Figure 1.**
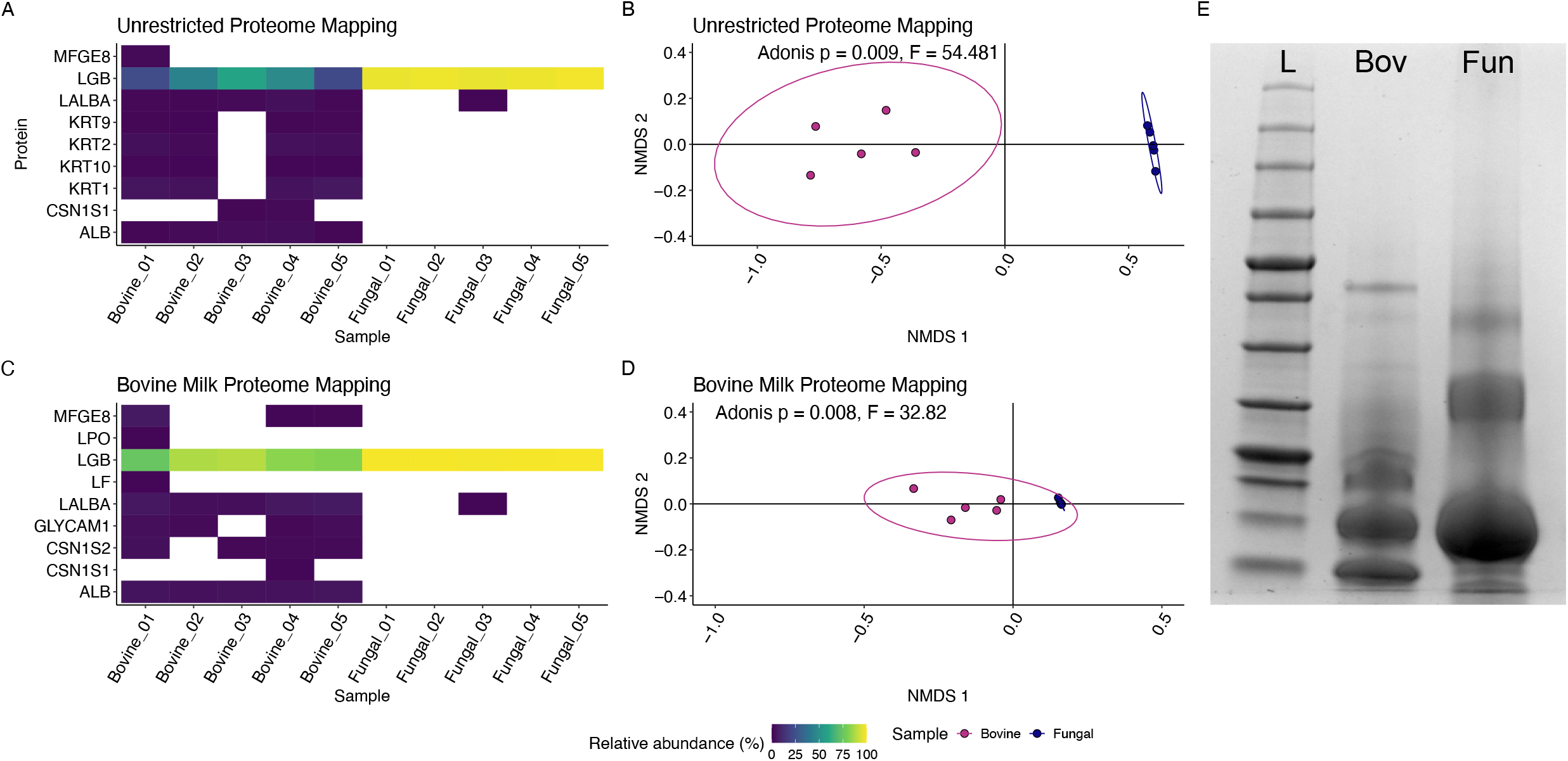
Characterization of protein samples by mass spectrometry identified similarities and differences. (A) Peptides mapped to an unrestricted protein database and were present at greater than 1% abundance were plotted and are colored by the relative abundance of the proteins. These results were also compared using a Bray Curtis distance showing significant differences between the samples (B; ADONIS, p = 0.008), here visualized here using nonmetric multidimensional scaling (NMDS). (C) In parallel, mapping peptides to a milk-specific protein database and (D) compared using the Bray Curtis distance showing significant differences as well (ADONIS, p = 0.009), visualized by NMDS. Proteins used to digest input protein samples were also detected, but were omitted from the analysis (porcine trypsin and a serine protease from *Achromabacter*). These findings were in agreement with SDS-PAGE of the same protein samples (E), showing a ladder (L), bovine whey (Bov) and fungal-derived whey (Fun).

In stark contrast to the bovine whey protein isolate, both the unrestricted protein database and the milk protein database for peptide mapping identified the most abundant protein as β-lactoglobulin in the yeast-derived whey protein isolate (>98% relative abundance; Figure 1, Table 1, S1). These findings were in agreement with analysis of these samples by SDS-PAGE, which found that in both sample types, β-lactoglobulin was the most common protein in both samples, with significantly less protein diversity in the yeast-derived whey protein sample (Figure 1C). While β-lactoglobulin was the most abundant protein in both samples, we compared the proteomes of the two samples using a Bray Curtis dissimilarity metric and compared the protein composition of these samples using an Adonis test. Irrespective of the mapping library used for the peptide mapping, we found that the composition of the protein samples were significantly different (p < 0.001; Figure 2) only when excluding β-lactoglobulin from the analysis. In the case of both protein samples, SDS-PAGE supported the findings by mass spectrometry (Figure 1C).

**Figure 2.**
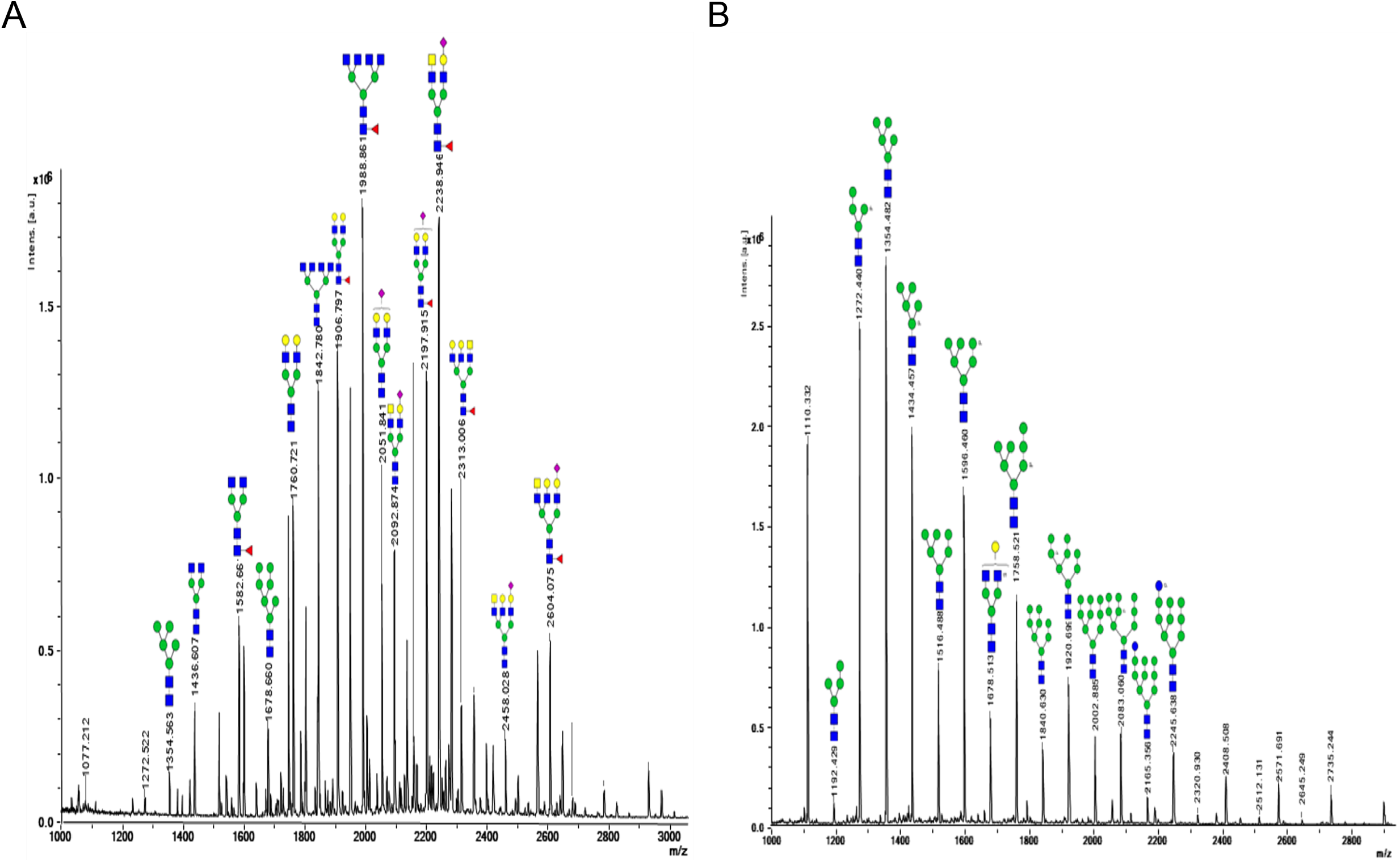
Identification of *N*-glycans identified in bovine whey protein and yeast-derived whey protein. Proteins released from bovine whey protein isolate and yeast-derived whey protein by PNGase-F were identified by mass spectrometry. (A) The most abundant *N*-glycans from bovine whey identified by mass spectra are shown. These glycans were characterized as being high mannose, complex, and hybrid, and fucosylated complex and hybrid *N*-glycans were identified. Six of the fifteen most abundant *N*-glycans in bovine whey protein were identified as sialylated, while the remaining nine were considered neutral *N*-glycans. (B) In contrast to the bovine whey, the fungal-derived *N*-glycans all structures were identified as neutral, high mannose *N*-glycans without fucosylation.

### 3.2 N-glycan analysis and comparison to bovine whey

*N*-glycans were released from protein samples using enzymatic deglycosylation with PNGase F and analyzed by matrix-assisted laser desorption/ionization time-of-flight coupled with mass spectrometry (MALDI-TOF MS) as described in the Methods section. The resulting mass spectra (Figure 3) were used to identify distinct glycan structures, which included 78 structures from the bovine whey protein sample and 22 total structures from the yeast-derived whey proteins. *N-*glycans are broadly characterized to be either high mannose, hybrid, or complex *N*-glycans, depending on their architecture and modification and can be further distinguished by whether they are decorated with sialic acid, yielding neutral or acidic *N*-glycan structures. Here, we found that the *N*-glycome of the bovine whey protein isolate contained 78 distinct structures, which included 9 high mannose, neutral *N*-glycans; 4 neutral hybrid *N*-glycans; 2 acidic hybrid *N*-glycans; 31 neutral complex *N*-glycans; and 32 acidic complex *N*-glycans. In contrast, the *N*-glycome of the yeast-produced whey protein contained 22 structures, with 16 neutral high mannose *N*-glycans and 6 neutral hybrid *N*-glycans. Of these glycans, 10 structures were shared between the two sample types, while 12 were unique to the yeast-derived whey and 67 were unique to bovine whey.

**Figure 3.**
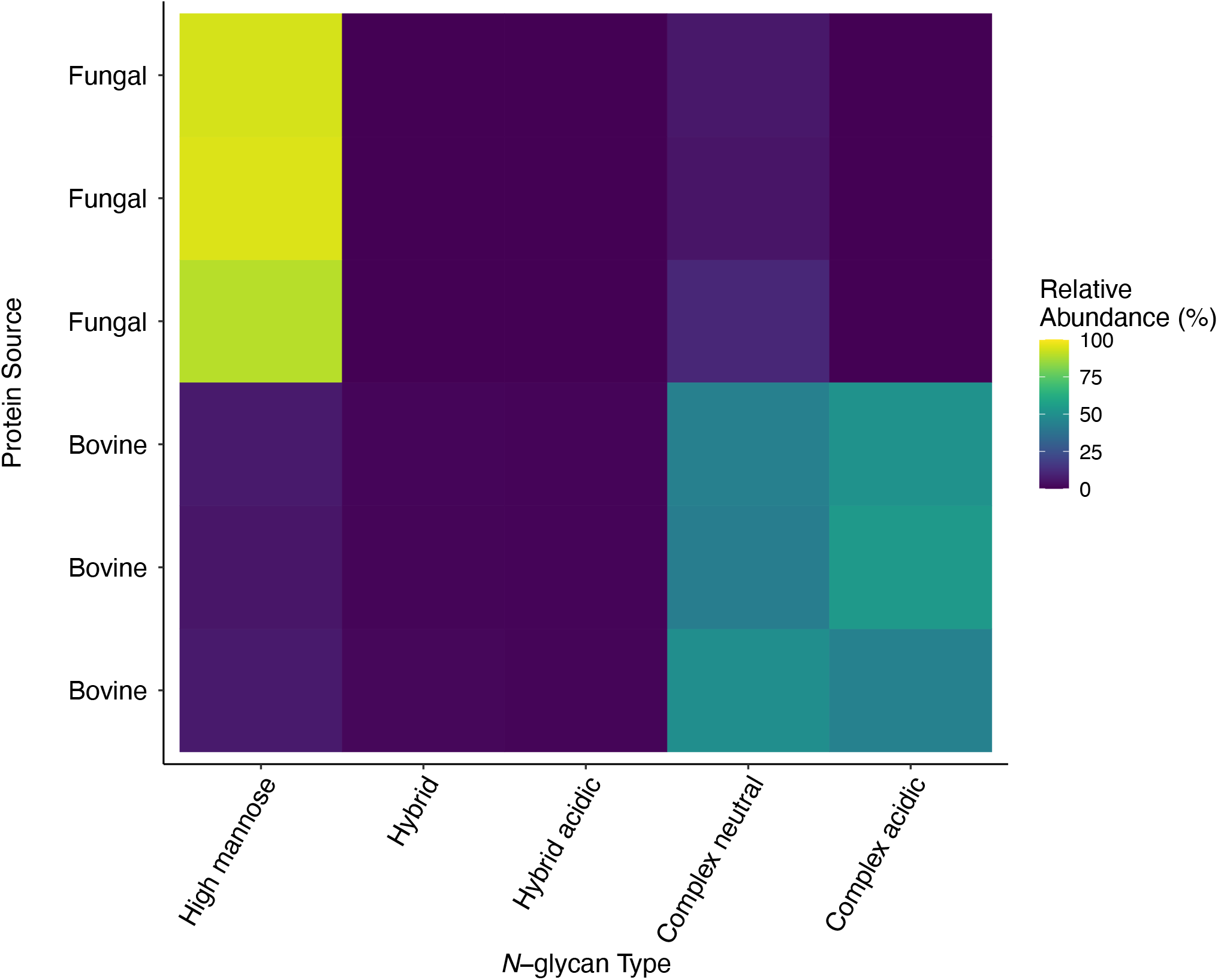
*N*-glycan class relative abundance among bovine and protein samples. *N*-glycans released from bovine whey protein isolate and yeast-derived whey protein samples by PNGase-F were identified by mass spectrometry and classified by composition to either neutral or acidic (containing sialic acid) and high mannose, complex, or hybrid structure. The relative abundance of each of these classes of *N*-glycans are colored by abundance across the tested samples.

The composition and characterization of these *N*-glycans are presented in Table 2. The relative abundance of neutral high mannose *N*-glycans as a fraction of the total glycome was higher in the yeast-produced whey protein sample, while there were more neutral and acidic complex *N-*glycans in the bovine whey protein glycome (Figure 3, 4).

**Figure 4.**
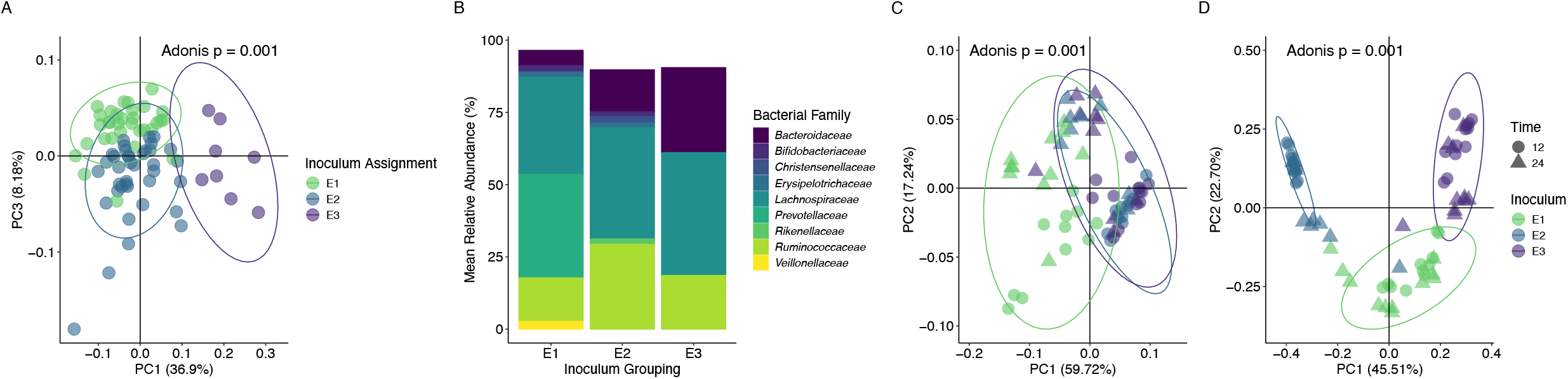
Microbial communities remain distinct at 12 and 24 hours. (A) Microbiome compositions of input fecal samples (B) Family-level stack plots of the starting inocula demonstrate distinct microbiome compositional state (Adonis p = 0.001), and after fermentation, distinct compositions remain at 12- and 24-hours (p = 0.001) when assessed by (C) weighted UniFrac and (D) Bray-Curtis dissimilarity measures. Bacterial families with a mean relative abundance less than 1.5% are omitted from the stacked bar plot.

### 3.3 Microbial communities remain distinct at 12 and 24 hours of in vitro fermentation

We first evaluated the fidelity of the *in vitro* system to faithfully recapitulate the distinct compositions detected in the input fecal samples. First, 16S rRNA gene sequencing identified three significantly distinct microbial community clusters (ADONIS, p = 0.001, Figure 4A) among fecal samples. We then followed up with pairwise PERMANOVA comparisons and observed significant differences between each microbial community cluster (FDR-adjusted p = 0.001). These clusters were identified as previously described (27), and six samples from each cluster were randomly selected for pooling to create the three microbial communities tested here (Figure 4B) and in our previous work (30). First, we tested whether these three representative microbial communities remained distinct during *in vitro* fermentation or whether the *in vitro* method collapsed community diversity to common features. We compared the three distinct communities grown on our soluble starch control at 12- and 24-hours post-inoculation. Using the weighted UniFrac distance metric, we found that the three distinct communities from the inocula remained distinct at 12- and 24-hours (ADONIS, p = 0.001, Figure 4C), which was corroborated by the non-phylogenetic Bray Curtis distance metric at 12- and 24-hours (ADONIS, p = 0.001, Figure 4D). Post-hoc PERMANOVA comparisons using the Bray Curtis metric showed that each community remained distinct at 12- and 24-hours (FDR-adjusted P < 0.05).

### 3.4 Impact of glycoprotein source on human gut microbial communities

To test the impact of our treatments within each representative microbial community, we compared samples exposed to our control of soluble starch, bovine, and fungal whey protein using the weighted UniFrac and Bray Curtis distance metrices. We found that across the three groups, and at both 12- and 24-hours, there were significant differences between the communities (ADONIS, p < 0.05) when compared using the weighted UniFrac and Bray Curtis distance metric (Figure 4C and 4D, respectively). Weighted UniFrac pairwise PERMANOVA comparisons did not find individual significant differences between E2 and E3 communities with bovine glycoproteins at 12 hours and between all three inocula grown on fungal or whey glycoproteins at 24 hours, after corrections for multiple comparisons (FDR-adjusted p > 0.05). In contrast, pairwise PERMANOVA comparisons using the Bray Curtis distance metric found significant differences between inocula on the substrates at both 12 and 24 hours (FDR-adjusted p < 0.05) except for E2 and E3 at 24 hours on bovine whey and between all three inocula at 24 hours on fungal whey (FDR-adjusted p > 0.05). Additional pairwise comparisons within microbial communities across substrates at 12 or 24 hours found that E1 grown on bovine whey protein was significantly different in terms of community composition at 24 hours when compared to the control and fungal whey protein (FDR-adjusted p < 0.05) when assessed by the Bray Curtis distance metric, while E2 grown with fungal whey protein was significantly different from the bovine and control groups at 12 hours. Inocula E3, by contrast, showed no significant differences after correction for multiple comparisons (FDR-adjusted p > 0.05).

To test whether distinct *N*-glycan structures affect diversity among these *in vitro* microbial communities, we compared the Shannon diversity of each microbial community exposed to bovine and fungal whey glycoproteins. At 12 hours, communities incubated with bovine whey protein were significantly more diverse than the control (FDR-adjusted P < 0.05, Figure 5A-C), but other comparisons were not significantly different (FDR-adjusted p > 0.05). At 24 hours, the communities incubated with bovine whey protein were significantly more diverse than the control (FDR-adjusted p < 0.001) and the communities incubated with the fungal whey protein (FDR-adjusted p < 0.05, Figure 5D-F). While the communities incubated with fungal whey protein trended higher in terms of diversity than the control, these differences were not significant at 12 or 24 hours (Figure A-F).

**Figure 5.**
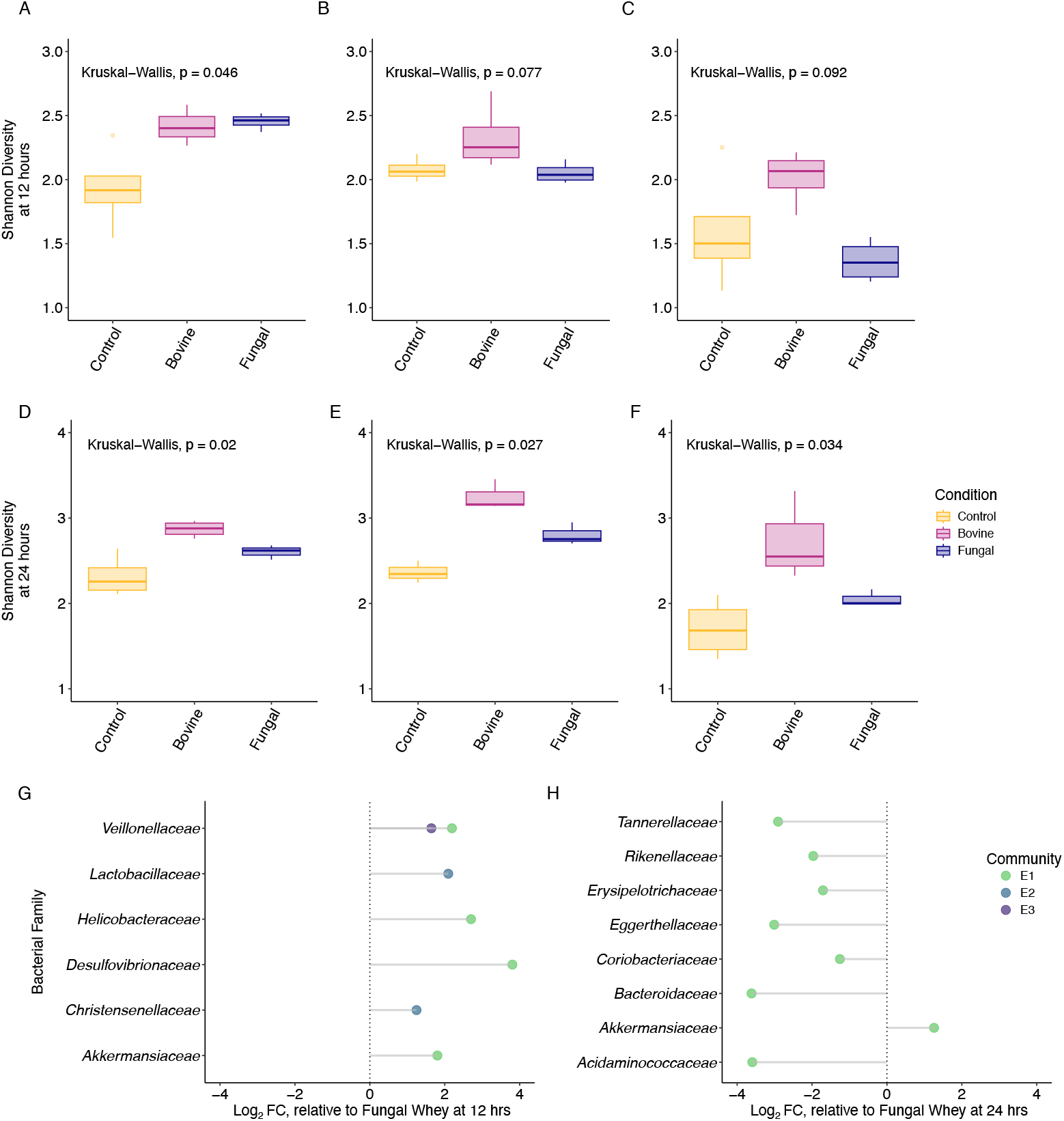
Alpha diversity and family-level taxa differ across microbial communities and substrates. Microbial communities (columns) show distinct responses to substrates (conditions) in terms of Shannon diversity at 12 and 24 hours (A-C, D-F, respectively), and ANCOM-BC determined glycoprotein sources enrich taxa at the family level in a substrate and community specific manner at 12- and 24-hours (G and H, respectively).

Finally, we performed Analysis of Compositions of Microbiomes with Bias Correction (ANCOM-BC) to identify differentially abundant taxa at the family-level within each microbial community in response to either bovine or fungal whey glycoproteins at 12- and 24-hours fermentation. All three starting microbial communities displayed significant enrichment in family-level taxa at 12 hours fermentation when grown on bovine whey relative to fungal whey (q < 0.05; Figure 5G), suggesting that the differences in *N-* glycan architecture between glycoprotein sources were driving family-level enrichment. Within microbial community E1 at 12 hours fermentation, bovine whey significantly enriched for *Veillonellaceae, Helicobacteraceae, Desulfovibrionaceae*, and *Akkermansiaceae* in relation to the fungal whey. Within microbial community E2 at 12 hours fermentation, bovine whey significantly enriched for *Lactobacteraceae* and *Christensenellaceae*. Lastly, within microbial community E3 at 12 hours, the only family that was differentially abundant was *Veillonellaceae*, which was significantly enriched on bovine relative to the fungal whey. Interestingly, microbial communities E2 and E3 did not contain any significant differences in family enrichment at 24 hours of fermentation, however, microbial community E1 contained eight families that significantly differed in abundance between the glycoprotein substrates. Within microbial community E1 at 24 hours fermentation, bovine whey only significantly enriched for *Akkermansiaceae*, while fungal whey enriched for *Tannerellaceae, Rikenellaceae, Erysipelotrichaceae, Eggerthellaceae, Coriobacteriaceae, Bacteroidaceae*, and *Acidaminococcaceae* (Figure 5H). Overall, no family was significantly enriched across all three microbial communities on either glycoprotein at 12- and 24-hours fermentation, suggesting community-specific responses to the glycoproteins. Interestingly, *Akkermansiaceae* within microbial community E1 was the only family significantly enriched across 12- and 24-hours fermentation. No other family remained significantly enriched across both timepoints on either glycoprotein substrate (Supplementary Figure 2).

## 4. Discussion

Advances in biosynthetic technologies have rapidly reshaped the development of food ingredients, with an aim to meet consumer perceptions about animal agriculture and a desire for improved efficiency in food production (1). However, despite technological advancements, these novel technologies cannot yet fully recapitulate the biological systems which have historically produced these food ingredients, even if the synthesis of individual constituents is technologically feasible (11). Here, we evaluated the composition of a novel bovine whey protein ingredient that is used to replace traditional bovine milk protein in foods in terms of its proteomic composition and the post-translational glycan modifications of these proteins made using distinct production sources. Using mass spectrometry, we characterized the proteomic composition and *N*-glycome of these two distinct protein samples.

While bovine-derived whey and the synthetic whey ingredient are both principally composed of β-lactoglobulin, the synthetic whey protein was almost entirely β-lactoglobulin, notably higher in relative abundance compared to bovine-derived whey. The remaining proteins detected in the yeast-derived whey could have originated from the production organism. In contrast, the bovine whey protein isolate was composed of more proteins, present at greater than 1% of the total composition, and exhibited greater total diversity relative to the yeast-derived whey. Consistent with previous work on bovine whey, these other proteins included α-lactalbumin, albumin, and casein S1 (17,40). While these protein samples were significantly different in terms of their composition (Figure 2), the overwhelming predominance of β-lactoglobulin in both samples (83% vs. 98%) suggests that the functional properties of both ingredients may be largely comparable (17), though we did not examine the effects of processing on glycation, nor the functional effects this may have, in the present work (12).

However, in stark contrast to the protein content, the *N*-glycome of these whey protein samples was remarkably distinct. Although β-lactoglobulin, not considered an *N*-glycosylated protein (41), was the predominant protein in the yeast-derived whey protein sample, we detected a wide variety of *N*-glycans in the yeast-derived whey protein sample that included a large number of unique structures absent in the bovine whey protein (Figure 3, Table 2). All *N*-glycans synthesized within eukaryotic organisms share a core structure of two 4Glc*N*Acβ1 sugars and a single mannose stemming into two branched mannose monomers (42). Depending on the organism, the final *N*-glycan is extended from this core through distinct biosynthetic pathways, resulting in a variety of *N*-glycan structures that reflect the machinery in the host cell. For example, yeast-derived *N*-glycans are structurally distinct from the hybrid or complex *N*-glycans are often reported for bovine milk protein *N*-glycans and the heterologous expression of mammalian proteins in yeast requires significant modification to the yeast glycosylation machinery to mimic the *N*-glycosylation of mammals (43,44). Further, it is important to note that *N*-glycans affect multiple characteristics of glycoproteins such as confirmation, solubility, antigenicity, activity, and recognition by glycan binding proteins (13,42,45,46). While β-lactoglobulin is not thought to be *N*-glycosylated (41), other synthetic food proteins with bioactive function that are *N-*glycosylated are likely to suffer functional deficits if glycosylation is not addressed in their manufacture.

Finally, there is existing evidence that *N*-glycans can serve as substrates for members of the gut microbiome, and specific adaptations for *N*-glycan utilization among gut microbes has also been described (14,15,47). We found that despite similar protein compositions between bovine and fungal whey glycoproteins, fermentation of these glycoproteins after 12 hours revealed family-level differential abundance across all three microbial communities’ representative of human gut microbiomes. Given our findings relating to *N-*glycan composition, we hypothesize that the differential abundance between glycoproteins at 12 hours were driven by the distinct glycosylation patterns observed, as well as the glycan fermentation capacity of each microbial community. However, these assays were performed *in vitro*, which is a limitation of the present study. Further testing will be necessary to characterize how *N-* glycan utilization differs among gut microbiome compositions, whether distinct genetic repertoires are found among these microbiome compositional types, and to identify which functional pathways are responsible for these differences.

In conclusion, we found that while the protein composition of a novel yeast-derived whey protein ingredient is distinct to bovine-derived whey protein isolate, the most distinguishing characteristic was the post-translational modifications to the respective proteins, depending on their origin. Thus, while we examined two commercially available dietary protein ingredients with comparable functionality, we found deeper distinctions between them that may affect bioactive functionality in other systems that use similar biosynthetic machinery.

## 5. Data availability

Proteomics and MALDI-TOF MS spectra are available in the Dryad data repository (doi:10.5061/dryad.hmgqnk9qz). 16S rRNA amplicon sequencing data is deposited in the NCBI SRA under BioProject PRJNA984714.

## 6. Acknowledgements

This work was made possible by funding from the University of Nevada, Reno Department of Nutrition, the College of Agriculture, Biotechnology, and Natural Resources, the Nevada Agriculture Experimental Station, and the University of Nevada, Reno Office of the Vice President for Research and Innovation. This work is also supported by a New Investigator Seed Grant from the USDA National Institute of Food and Agriculture, #13385133 and by funding from the National Institute of General Medical Sciences (GM103440 and GM104944) from the National Institutes of Health. The authors thank the Mitch Hitchcock Ph.D. Nevada Proteomics Center (RRID:SCR_017761) and the Nevada Bioinformatics Center (RRID:SCR_017802) for their technical support as well as the Molecular Research Core Facility at Idaho State University, RRID:SCR_012598. The authors also thank Dr. David C Dallas (Oregon State University; http://www.dallaslab.org) for providing the bovine milk specific protein database used in the proteomic analysis.

## 7. Disclosures

The authors have no conflicts of interest to disclose.

## 8. Figure Legends

**Supplementary Figure 1.**
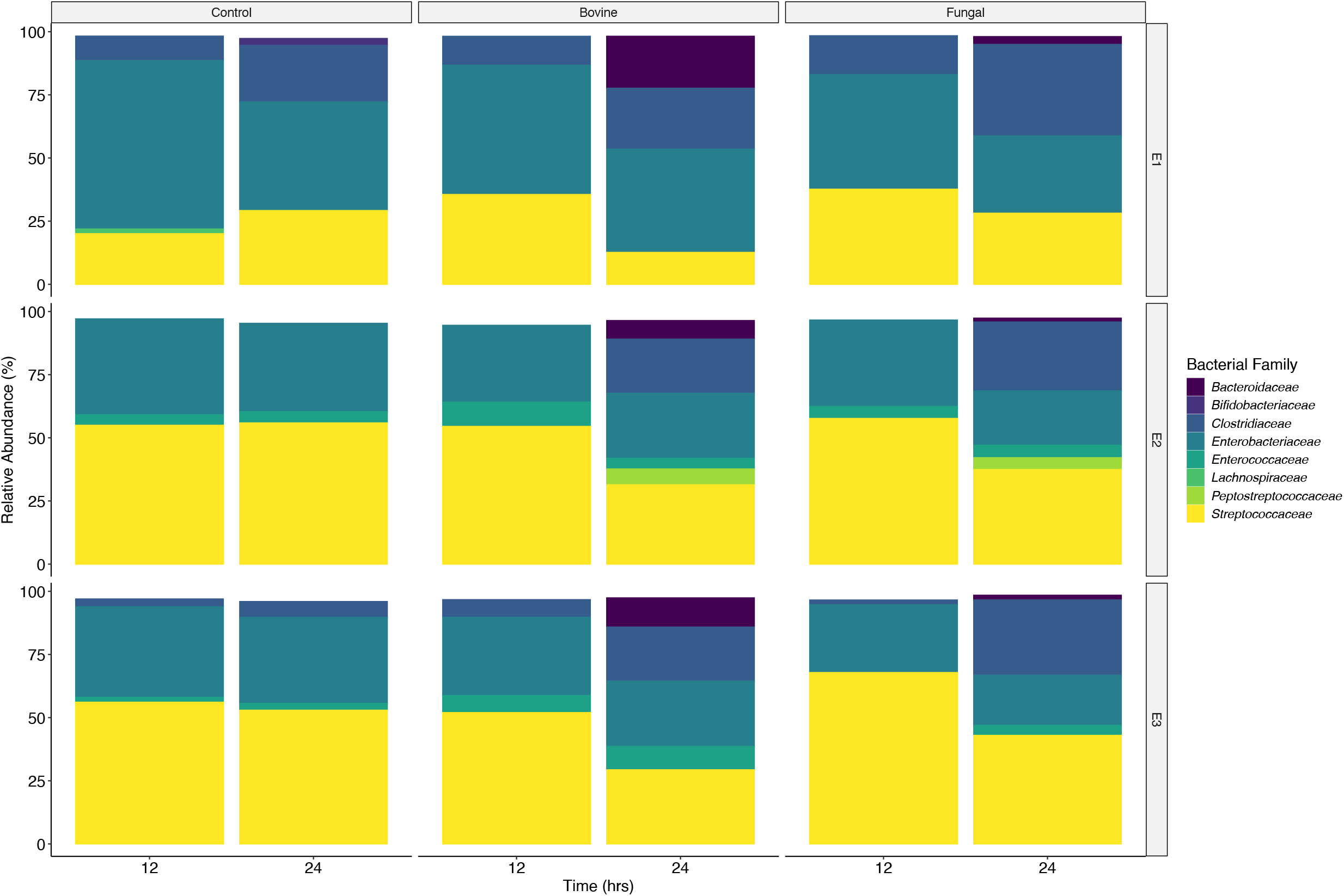
Stack bar plots showing the relative abundance of family-level taxa within each microbial community after fermentation of respective glycoprotein or starch substrates at 12- and 24-hours. Bacterial families with a mean relative abundance less than 1.5% are omitted.

## 9. Table legends

**Table 1.** Relative abundance of proteins mapped using a milk-protein specific database.

**Table 2.** *N*-glycans identified across whey protein isolate samples.

**Table S1.** Relative abundance of proteins mapped using an unrestricted proteome database (Spectronaut).

## Notes

### Competing Interest Statement

The authors have declared no competing interest.

## References

1. Hartmann C, Siegrist M. Consumer perception and behaviour regarding sustainable protein consumption: A systematic review. Trends Food Sci Technol (2017) 61:11–25. doi: 10.1016/j.tifs.2016.12.006

2. McClements DJ, Newman E, McClements IF. Plant-based Milks: A Review of the Science Underpinning Their Design, Fabrication, and Performance. Compr Rev Food Sci Food Saf (2019) 18:2047–2067. doi: 10.1111/1541-4337.12505

3. Vanga SK, Raghavan V. How well do plant based alternatives fare nutritionally compared to cow’s milk? J Food Sci Technol (2018) 55:10–20. doi: 10.1007/s13197-017-2915-y

4. Merritt RJ, Fleet SE, Fifi A, Jump C, Schwartz S, Sentongo T, Duro D, Rudolph J, Turner J, NASPGHAN Committee on Nutrition. North American Society for Pediatric Gastroenterology, Hepatology, and Nutrition Position Paper: Plant-based Milks. J Pediatr Gastroenterol Nutr (2020) 71:276–281. doi: 10.1097/MPG.0000000000002799

5. Paul AA, Kumar S, Kumar V, Sharma R. Milk Analog: Plant based alternatives to conventional milk, production, potential and health concerns. Crit Rev Food Sci Nutr (2020) 60:3005–3023. doi: 10.1080/10408398.2019.1674243

6. Bych K, Mikš MH, Johanson T, Hederos MJ, Vigsnæs LK, Becker P. Production of HMOs using microbial hosts - from cell engineering to large scale production. Curr Opin Biotechnol (2019) 56:130–137. doi: 10.1016/j.copbio.2018.11.003

7. Mendes A, Reis A, Vasconcelos R, Guerra P, Lopes da Silva T. Crypthecodinium cohnii with emphasis on DHA production: a review. J Appl Phycol (2009) 21:199–214. doi: 10.1007/s10811-008-9351-3

8. Chen L, Guttieres D, Koenigsberg A, Barone PW, Sinskey AJ, Springs SL. Large-scale cultured meat production: Trends, challenges and promising biomanufacturing technologies. Biomaterials (2022) 280:121274. doi: 10.1016/j.biomaterials.2021.121274

9. Chang MCY, Keasling JD. Production of isoprenoid pharmaceuticals by engineered microbes. Nat Chem Biol (2006) 2:674–681. doi: 10.1038/nchembio836

10. Ro D-K, Paradise EM, Ouellet M, Fisher KJ, Newman KL, Ndungu JM, Ho KA, Eachus RA, Ham TS, Kirby J, et al. Production of the antimalarial drug precursor artemisinic acid in engineered yeast. Nature (2006) 440:940–943. doi: 10.1038/nature04640

11. Hettinga K, Bijl E. Can recombinant milk proteins replace those produced by animals? Curr Opin Biotechnol (2022) 75:102690. doi: 10.1016/j.copbio.2022.102690

12. Chobert J-M, Gaudin J-C, Dalgalarrondo M, Haertlé T. Impact of Maillard type glycation on properties of beta-lactoglobulin. Biotechnol Adv (2006) 24:629–632. doi: 10.1016/j.biotechadv.2006.07.004

13. Barboza M, Pinzon J, Wickramasinghe S, Froehlich JW, Moeller I, Smilowitz JT, Ruhaak LR, Huang J, Lönnerdal B, German JB, et al. Glycosylation of Human Milk Lactoferrin Exhibits Dynamic Changes During Early Lactation Enhancing Its Role in Pathogenic Bacteria-Host Interactions *. Mol Cell Proteomics (2012) 11: doi: 10.1074/mcp.M111.015248

14. Karav S, Casaburi G, Arslan A, Kaplan M, Sucu B, Frese S. N-glycans from human milk glycoproteins are selectively released by an infant gut symbiont in vivo. J Funct Foods (2019) 61:103485. doi: 10.1016/j.jff.2019.103485

15. Karav S, Le Parc A, Leite Nobrega de Moura Bell JM, Frese SA, Kirmiz N, Block DE, Barile D, Mills DA. Oligosaccharides Released from Milk Glycoproteins Are Selective Growth Substrates for Infant-Associated Bifidobacteria. Appl Environ Microbiol (2016) 82:3622–3630. doi: 10.1128/AEM.00547-16

16. Oftedal OT. The evolution of milk secretion and its ancient origins. animal (2012) 6:355–368. doi: 10.1017/S1751731111001935

17. Kelly P, Woonton BW, Smithers GW. “Improving the sensory quality, shelf-life and functionality of milk.,” In: Paquin P, editor. Functional and Speciality Beverage Technology. Woodhead Publishing Series in Food Science, Technology and Nutrition. Woodhead Publishing (2009). p. 170–231 doi: 10.1533/9781845695569.2.170

18. Kayili HM, Salih B. N-glycan Profiling of Glycoproteins by Hydrophilic Interaction Liquid Chromatography with Fluorescence and Mass Spectrometric Detection. J Vis Exp JoVE (2021) doi: 10.3791/62751

19. Frese SA, Hutton AA, Contreras LN, Shaw CA, Palumbo MC, Casaburi G, Xu G, Davis JCC, Lebrilla CB, Henrick BM, et al. Persistence of Supplemented Bifidobacterium longum subsp. infantis EVC001 in Breastfed Infants. mSphere (2017) 2: doi: 10.1128/mSphere.00501-17

20. Kozich JJ, Westcott SL, Baxter NT, Highlander SK, Schloss PD. Development of a Dual-Index Sequencing Strategy and Curation Pipeline for Analyzing Amplicon Sequence Data on the MiSeq Illumina Sequencing Platform. Appl Environ Microbiol (2013) 79:5112–5120. doi: 10.1128/AEM.01043-13

21. Parada AE, Needham DM, Fuhrman JA. Every base matters: assessing small subunit rRNA primers for marine microbiomes with mock communities, time series and global field samples. Environ Microbiol (2016) 18:1403–1414. doi: 10.1111/1462-2920.13023

22. Apprill A, McNally S, Parsons R, Weber L. Minor revision to V4 region SSU rRNA 806R gene primer greatly increases detection of SAR11 bacterioplankton. Aquat Microb Ecol (2015) 75:129–137. doi: 10.3354/ame01753

23. Bolyen E, Rideout JR, Dillon MR, Bokulich NA, Abnet CC, Al-Ghalith GA, Alexander H, Alm EJ, Arumugam M, Asnicar F, et al. Reproducible, interactive, scalable and extensible microbiome data science using QIIME 2. Nat Biotechnol (2019) 37:852–857. doi: 10.1038/s41587-019-0209-9

24. Callahan BJ, McMurdie PJ, Rosen MJ, Han AW, Johnson AJA, Holmes SP. DADA2: High-resolution sample inference from Illumina amplicon data. Nat Methods (2016) 13:581–583. doi: 10.1038/nmeth.3869

25. Price MN, Dehal PS, Arkin AP. FastTree 2--approximately maximum-likelihood trees for large alignments. PloS One (2010) 5:e9490. doi: 10.1371/journal.pone.0009490

26. Pruesse E, Quast C, Knittel K, Fuchs BM, Ludwig W, Peplies J, Glöckner FO. SILVA: a comprehensive online resource for quality checked and aligned ribosomal RNA sequence data compatible with ARB. Nucleic Acids Res (2007) 35:7188–7196. doi: 10.1093/nar/gkm864

27. Arumugam M, Raes J, Pelletier E, Le Paslier D, Yamada T, Mende DR, Fernandes GR, Tap J, Bruls T, Batto J-M, et al. Enterotypes of the human gut microbiome. Nature (2011) 473:174–180. doi: 10.1038/nature09944

28. Mandal S, Van Treuren W, White RA, Eggesbø M, Knight R, Peddada SD. Analysis of composition of microbiomes: a novel method for studying microbial composition. Microb Ecol Health Dis (2015) 26:27663. doi: 10.3402/mehd.v26.27663

29. Aranda-Díaz A, Ng KM, Thomsen T, Real-Ramírez I, Dahan D, Dittmar S, Gonzalez CG, Chavez T, Vasquez KS, Nguyen TH, et al. Establishment and characterization of stable, diverse, fecal-derived in vitro microbial communities that model the intestinal microbiota. Cell Host Microbe (2022) 30:260-272.e5. doi: 10.1016/j.chom.2021.12.008

30. Flores Martinez KE, Bloszies CS, Bolino MJ, Henrick BM, Frese SA. Hemp hull fiber and two constituent compounds, N-trans-caffeoyltyramine and N-trans-feruloyltyramine, shape the human gut microbiome in vitro. Food Chem X (2024) 23:101611. doi: 10.1016/j.fochx.2024.101611

31. R: A language and environment for statistical computing. (2022)

32. Kruskal WH, Wallis WA. Use of Ranks in One-Criterion Variance Analysis. J Am Stat Assoc (1952) 47:583–621. doi: 10.1080/01621459.1952.10483441

33. Kassambara A. ggpubr: “ggplot2” Based Publication Ready Plots. (2022) https://CRAN.R-project.org/package=ggpubr [Accessed January 9, 2023]

34. Kassambara A. rstatix: Pipe-Friendly Framework for Basic Statistical Tests. (2022) https://CRAN.R-project.org/package=rstatix [Accessed January 9, 2023]

35. Bray JR, Curtis JT. An Ordination of the Upland Forest Communities of Southern Wisconsin. Ecol Monogr (1957) 27:326–349. doi: 10.2307/1942268

36. Oksanen J, Simpson GL, Blanchet FG, Kindt R, Legendre P, Minchin PR, O’Hara RB, Solymos P, Stevens MHH, Szoecs E, et al. vegan: Community Ecology Package. (2022) https://CRAN.R-project.org/package=vegan [Accessed January 9, 2023]

37. Wickham H, RStudio. tidyverse: Easily Install and Load the “Tidyverse.” (2022) https://CRAN.R-project.org/package=tidyverse [Accessed January 9, 2023]

38. Wickham H, Chang W, Henry L, Pedersen TL, Takahashi K, Wilke C, Woo K, Yutani H, Dunnington D, RStudio. ggplot2: Create Elegant Data Visualisations Using the Grammar of Graphics. (2022) https://CRAN.R-project.org/package=ggplot2 [Accessed January 9, 2023]

39. Garnier S, Ross N, Rudis B, Sciaini M, Camargo AP, Scherer C. viridis: Colorblind-Friendly Color Maps for R. (2021) https://CRAN.R-project.org/package=viridis [Accessed January 9, 2023]

40. Kinsella JE, Morr CV. Milk proteins: Physicochemical and functional properties. C R C Crit Rev Food Sci Nutr (1984) 21:197–262. doi: 10.1080/10408398409527401

41. Valk-Weeber RL, Nichols K, Dijkhuizen L, Bijl E, van Leeuwen SS. Variations in N-linked glycosylation of glycosylation-dependent cell adhesion molecule 1 (GlyCAM-1) whey protein: Intercow differences and dietary effects. J Dairy Sci (2021) 104:5056–5068. doi: 10.3168/jds.2020-19297

42. Stanley P, Taniguchi N, Aebi M. “N-Glycans.,” In: Varki A, Cummings RD, Esko JD, Stanley P, Hart GW, Aebi M, Darvill AG, Kinoshita T, Packer NH, Prestegard JH, et al., editors. Essentials of Glycobiology. Cold Spring Harbor (NY): Cold Spring Harbor Laboratory Press (2015) http://www.ncbi.nlm.nih.gov/books/NBK453020/ [Accessed January 9, 2023]

43. Hamilton SR, Bobrowicz P, Bobrowicz B, Davidson RC, Li H, Mitchell T, Nett JH, Rausch S, Stadheim TA, Wischnewski H, et al. Production of complex human glycoproteins in yeast. Science (2003) 301:1244–1246. doi: 10.1126/science.1088166

44. Nwosu CC, Aldredge DL, Lee H, Lerno LA, Zivkovic AM, German JB, Lebrilla CB. Comparison of the human and bovine milk N-glycome via high-performance microfluidic chip liquid chromatography and tandem mass spectrometry. J Proteome Res (2012) 11:2912–2924. doi: 10.1021/pr300008u

45. Lisowska E. The role of glycosylation in protein antigenic properties. Cell Mol Life Sci CMLS (2002) 59:445–455. doi: 10.1007/s00018-002-8437-3

46. Schwarz F, Aebi M. Mechanisms and principles of N-linked protein glycosylation. Curr Opin Struct Biol (2011) 21:576–582. doi: 10.1016/j.sbi.2011.08.005

47. Duman H, Kaplan M, Arslan A, Sahutoglu AS, Kayili HM, Frese SA, Karav S. Potential Applications of Endo-β-N-Acetylglucosaminidases From Bifidobacterium longum Subspecies infantis in Designing Value-Added, Next-Generation Infant Formulas. Front Nutr (2021) 8: doi: 10.3389/fnut.2021.646275

